# Computational identification and characterization of antigenic properties of Rv3899c of *Mycobacterium tuberculosis* and its interaction with Human leukocyte antigen (HLA)

**DOI:** 10.1101/2021.01.10.426101

**Authors:** Ritam Das, Kandasamy Eniyan, Urmi Bajpai

**Author notes:** **Address for communication:** Dr.Urmi Bajpai (Associate Professor) Department of Biomedical Science, Acharya Narendra Dev College (University of Delhi), Govindpuri, Kalkaji, New-Delhi-110019,India. Email id.

## Abstract

Tuberculosis (TB) is the second largest infectious disease that kills 1.2 million people annually worldwide. A rise in drug-resistant TB cases demands continued efforts towards the discovery & development of drugs and vaccines. In the recent past, though novel drugs have been added to the current TB regimen [1], research on new vaccine candidates needs a greater thrust. Secretory proteins of H37Rv are frequently studied for their antigenicity and their scope as protein subunit vaccines decrees further analysis. In this study, Rv3899c of H37Rv emerges as a potential vaccine candidate on its evaluation by several bioinformatics tools. It is a non-toxin, secretory protein with an ‘immunoglobulin-like’ fold which does not show similarity with a human protein. We found Rv3899c homologs in several mycobacterial species and its antigenic score (0.54) to compare well with the known immunogens such as ESAT-6 (0.56) and Rv1860 (0.52). Structural examination of Rv3899c predicted ten antigenic peptides, an accessibility profile of the antigenic determinants constituting B-cell epitope rich regions and a low Abundance of Antigenic Regions (AAR) value. Significantly, our study shows ESX-2 secretion system proteins and antigenic PE/PPE proteins of H37Rv as the interacting partners of Rv3899c. Further, molecular docking predicted Rv3899c to interact with human leukocyte antigen HLA-DRB1*04:01 through its antigenically conserved motif (RAAEQQRLQRIVDAVARQEPRISWAAGLRDDGTT). Interestingly, the binding affinity was observed to increase on citrullation of its Arg1 residue. Taken together, the computational characterization and predictive information suggest Rv3899c to be a promising TB vaccine candidate, which should be examined and validated experimentally.

## 1. Introduction

Tuberculosis (TB) is one of the top ten causes of death in the world and the death rate is more than that of the fatal viral diseases such as HIV/AIDS. *Mycobacterium tuberculosis,* the causative agent of TB is estimated to cause 1.2 million deaths annually worldwide (WHO, 2019) [2]. Presently, we are witnessing the emergence of multidrug-resistant (MDR) and extensively drug-resistant (XDR) strains of *Mycobacterium tuberculosis*. In the case of vaccines, an anti-TB vaccine (BCG) developed by Calmette and Guerin from *Mycobacterium bovis*has been used in several countries, however, the defensive potency of this is limited to young children with minimal effectiveness in adults [3, 4]. Hence, there is an urgent need for the improvement in TB diagnosis, formulation of new drug regimens and development of new anti-TB vaccines that can potentially assist in building up a promising strategy to tackle this global menace [5].Since BCG, which is the only TB vaccine approved in most countries, hasn’t been entirely effective, 13 new vaccines have been developed in the past decade and are currently being tested for their efficacy through clinical trials. However, the results have not been encouraging [3]. Due to the high risk involved in the administration of live (attenuated) vaccines against TB, mycobacterial protein and DNA vaccines are being currently studied as effective candidates [6].

Reverse vaccinology, an integral part of Immunoinformatics, has been gaining popularity to search for antigenic proteins as most of the infectious microorganisms cannot be cultured outside their hosts. It works by screening the proteome of the pathogen computationally in search of antigenic candidates that are preferably secretory or are membrane proteins, consists of B-cell or T-cell epitopes and show no similarity with a human protein [7]. In the past, numerous studies have validated the importance of secretory proteins of *M. tuberculosis* as potential immunogens. In one of the reports, the dose-dependent protective immune response was noted in mice when administered with the secretory proteins ESAT-6 and antigen 85B from *M. tuberculosis*[8]. While another study showed Rv1860 secretory protein can proliferate IFN-γ from human CD4+ and CD8+ T-cells and also successfully elicit a protective immune response in mice infected with a virulent strain of *M. tuberculosis*[9]. Hence, analysis of the secretory proteins of *M. tuberculosis* could help us identify and characterize the antigenic proteins that could elicit a protective immune response. In this paper, we have evaluated Rv3899c as an effective vaccine candidate using various bioinformatics tools. Earlier studies have identified Rv3899c as a secretory hypothetical protein and predicted it to be a putative immunogen [10, 11], however, a detailed examination of the protein and its potential as a vaccine candidate has not been done. Here,we have characterized this hypothetical protein *in silico* and found possible antigenic determinants and explored its potential as a subunit vaccine candidate (Figure 1). In terms of its secretory nature, antigenicity and the absence of a homolog in the human proteome, it was encouraging to find Rv3899c to be at par with the already known immunogens ESAT-6 and Rv1860 from *M.tuberculosis*. Additionally, it also contains several B-cell epitope rich regions and on a promising note, ‘motif 2’ determined in this study was found to be conserved over several mycobacterium species with an antigenicity score of 0.7 and to have a binding affinity for HLA-DRB1*0401 MHC class II molecule. Hence, the characterization carried out in this study strengthens Rv3899c as a TB vaccine candidate which should be experimentally verified.

**Figure 1:**
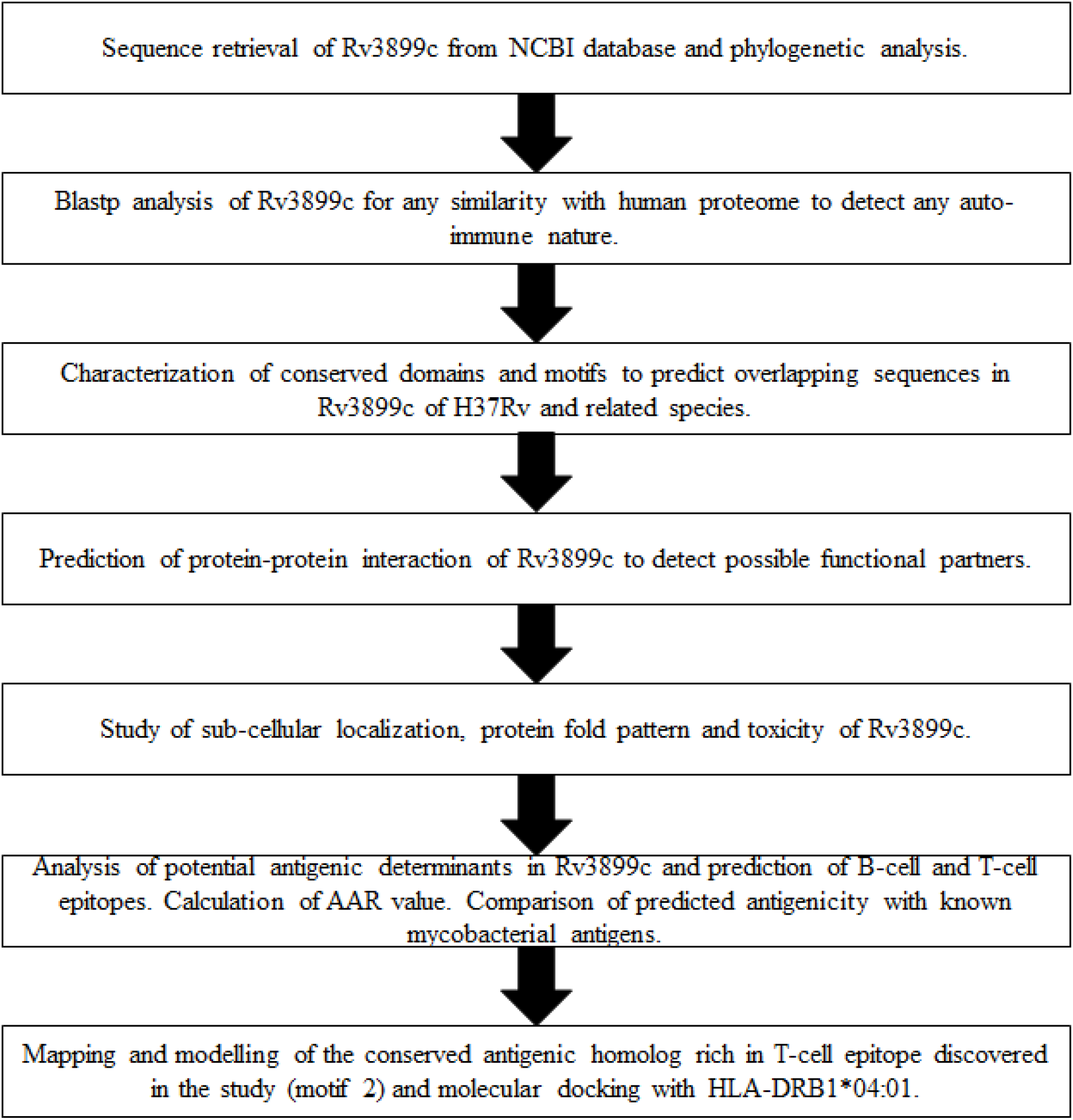
Overview of various bioinformatics analysis and steps involved in this study for the detection of potential antigenic determinants in Rv3899c.

## 2. Methods

### 2.1. Characterization of Rv3899c sequence from H37Rv

The nucleotide and protein sequences of Rv3889c were retrieved from NCBI database (https://www.ncbi.nlm.nih.gov/). Blastp suite (https://blast.ncbi.nlm.nih.gov/Blast.cgi) [12], ClustalW (https://www.genome.jp/tools-bin/clustalw) [13] and Phylogeny.fr (http://www.phylogeny.fr/) [14] were used for homology studies, multiple sequence alignment and to generate phylogenetic relationships of Rv3899c respectively.

### 2.2. Conserved domain and motif analysis in Rv3899c and related sequences

Conserved domains in Rv3889c were found using InterProScan 5 (https://www.ebi.ac.uk/) [15] and NCBI Conserved Domain Search tool (https://www.ncbi.nlm.nih.gov/Structure/cdd/wrpsb.cgi) [16]. To predict conserved overlapping motifs, Rv3889c and related protein sequences were evaluated using MEME suite (http://meme-suite.org/) [17] and their functions were predicted (threshold was 1.0) with the Motif Search tool (https://www.genome.jp/tools/motif/) [18].

### 2.3. In silico analysis of Rv3899c protein

To study the physio-chemical properties of the protein, the amino acid sequence was analysed through ProtParam tool (https://web.expasy.org/protparam/) ProtScale of ExPASy (https://web.expasy.org/protscale/) [19] was used to evaluate the hydrophobicity, polarity, flexibility and percentage accessibility profile of the protein. Potential sites of phosphorylation in the protein was analysed using NetPhos 3.1 server (http://www.cbs.dtu.dk/services/NetPhos/) [20] and the secondary structure prediction was done using PredictProtein Program (https://predictprotein.org/) [21].

### 2.4. Analysis of protein fold pattern and interaction network of Rv3899c

Recognition of the pattern of protein folding was done with PFP-FunDSeqE Server (http://www.csbio.sjtu.edu.cn/bioinf/PFP-FunDSeqE/) [22] and the identification of possible functional protein network was carried out using STRING 10.0 (https://string-db.org/) [23].

### 2.5. Prediction of subcellular localization and toxicity of Rv3899c

SignalP-5.0 (http://www.cbs.dtu.dk/services/SignalP/) [24] and TargetP-2.0 (http://www.cbs.dtu.dk/services/TargetP/) [25] were used to detect the presence of a signal peptide and SecretomeP-2.0 (http://www.cbs.dtu.dk/services/SecretomeP/) [26] was used to determine the secretory nature of the protein. Prediction of transmembrane helices in the protein was done using TMHMM Server v.2.0 (http://www.cbs.dtu.dk/services/TMHMM/) [27] and HMMTOP tool (http://www.enzim.hu/hmmtop/) [28]. Prediction of subcellular localization and toxicity of Rv3899c, based on various SVM modules, was done using PSLpred (http://crdd.osdd.net/raghava/pslpred/) [29] and BTXpred Server (http://crdd.osdd.net/raghava/btxpred/) [30], respectively.

### 2.6. Identification and characterization of antigenic determinants in Rv3899c

The antigenic potential of Rv3899c was determined using VirulentPred (http://203.92.44.117/virulent/) [31] and VaxiJen v2.0 (http://www.ddg-pharmfac.net/vaxijen/VaxiJen/VaxiJen.html) [32]. The bacterial model was selected and the protein was screened against a standard threshold of 0.4. The antigenic regions in Rv3889c were then mapped using Immunomedicine (http://imed.med.ucm.es/Tools/antigenic.pl) [33]. The regions predicted using this tool were again screened individually for their antigenicity. Prediction of B-cell epitopes were done using Bcepred (http://crdd.osdd.net/raghava/bcepred/) [34] and BepiPred-2.0 (http://www.cbs.dtu.dk/services/BepiPred/) [35]. The Abundance of Antigenic Regions (AAR) value of a protein is defined as the number of amino acids in between antigenic sequences predicted by software and low AAR value indicates higher antigenicity [36]. The AAR of Rv3899c was calculated as follows:

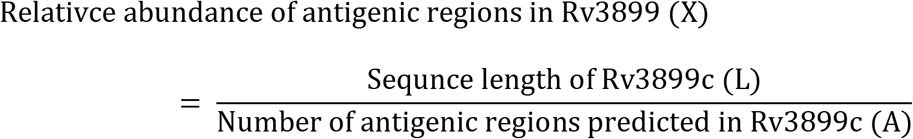

The prediction of T-cell epitopes for HLA-DRB1 04:01 in the antigenic regions in Rv3899c were done using SYFPEITHI (http://www.syfpeithi.de/) [37] and IEDB server (https://www.iedb.org/) [38]followed by evaluation using PepBind (https://yanglab.nankai.edu.cn/PepBind/) [39] to detect protein-peptide binding residues. PepBank (http://pepbank.mgh.harvard.edu/) was used to predict the presence of previously known antigenic peptides in Rv3899c through sequence text mining approach [40] and similarity analysis with human proteome was performed through Blastp suite (https://blast.ncbi.nlm.nih.gov/Blast.cgi) [12] to detect any autoimmune nature of Rv3899c. Also, a comparative analysis of Rv3899c with other antigens from H37Rv was performed on the basis of secretory and antigenicity score. Sequence of known mycobacterial antigens Rv1790 (PPE27) [41], Rv1860 (Alanine and proline-rich secreted protein Apa) [9], Ag85A (Diacylglycerol acyltransferase/mycolyltransferase), Ag85B (secreted antigen), Ag85c (Diacylglycerol acyltransferase/mycolyltransferase) [42], Rv3875 (esxA), Rv3874 (cfp-10) [43], Rv3418c (groS) [44], and Rv3621c (PPE65) [45] were analysed using the SecretomeP 2.0 and the VaxiJen v2.0 tool.

### 2.7. Mapping of antigenic homolog ‘motif 2’ in the secondary and tertiary structure of Rv3899c^184-410^

The crystal structure of a fragment of the protein (Rv3899c^184-410^) was obtained from Protein Data Bank (ID: 5IMU) (https://www.rcsb.org/) and the antigenic peptides were mapped on the 3D structure using PyMOL[46] and on the secondary structure too which was generated using PDBsum (http://www.ebi.ac.uk/thornton-srv/databases/cgi-bin/pdbsum/GetPage.pl?pdbcode=index.html) [47].

### 2.8. Molecular docking of ‘motif 2’ and human leukocyte antigen (HLA) complex (HLA-DRB1*04:01) using PatchDock

The 3D structure of the conserved antigenic ‘motif 2’ discovered in this study was constructed with the help of SwissModel server (https://swissmodel.expasy.org/) [48] using the crystal structure of Rv3899c^184-410^ (PDB ID: 5IMU) as a template. The structure obtained was further analyzed for its binding affinity for HLA-DRB1*04:01 [49] through shape complementarity molecular docking algorithm using the tool PatchDock (https://bioinfo3d.cs.tau.ac.il/PatchDock/patchdock.html) [50]. Blind docking of HLA-DRB1*04:01 as receptor and ‘motif 2’ as a ligand was performed and the complex with the lowest binding energy and high docking score was visually rescored using PyMOL.

Substitution of Arginine to Citrulline in the antigenic peptides has earlier been demonstrated to increase binding affinity for HLA-DRB1*04:01 [51]. To investigate whether citrullination has a similar effect in the case of ‘motif 2’ in our study, *in silico* citrullination of Arg1 (CITAAEQQRLQRIVDAVARQEPRISWAAGLRDDGTT) was carried out using PyTMs plugin of PyMOL [52] followed by molecular docking using PatchDock.

## 3. Results

### 3.1. Analysis of Rv3899c protein sequence

Sequence alignment and phylogenetic analysis revealed Rv3899c of *M. tuberculosis* H37Rv to share a 100% homology with Tat (twin-arginine translocation) pathway signal protein of *Mycobacterium sp.* 3/86Rv while 100% and 99.6% identity with *Mycobacterium tuberculosis* KZN 4207 and DUF5632 domain-containing protein *Mycobacterium shinjukuense,* respectively. Besides, there are DUF5631 and DUF5632 domain-containing proteins from several other mycobacterial species which also show 70% identity with Rv3899c (Figure S1).

### 3.2. Conserved domains and motifs predicted in Rv3899c

Domain analysis shows Rv3899c to have DUF5631 (Rv3899c^304-397^) and DUF5632 (Rv3899c^195-276^) Pfam alpha-helical domains with an unknown function found at the C-terminal region, known to be conserved across several mycobacterial species (Figure S2).

Rv3899c of H37Rv and its related homologs potentially have 5 conserved overlapping motifs designated as motif 1 to 5 respectively (Figure 2). Motif 1 (E-value: 7.6e-4733) located between amino acid positions ‘347-396’ has a width of 50 amino acid residue and consists of 3 conserved regions. The domain of unknown function 5631 (NCBI CDD 376071, E-value: 1.2e-21) was found to be conserved in the entire width of the sequence while region ‘10-35’ and ‘4-41’ of motif 1 were predicted to be a periplasmic component associated with an ABC type carbohydrate transport mechanism across membranes (NCBI CDD 224791, E-value 0.9) and to have a glycerol-3-phosphate acyltransferase role (NCBI CDD 237707, E-value 0.7), respectively. Motif 3 (E-value: 3.6e-4465) having a width of 50 amino acid residues (Rv3899c^231-280^) is predicted to have a conserved region consisting of Duffy like binding domain (NCBI CDD 368435, E-value 0.07). Characterization of motif 5 (E-value 1.9e-1450) present in the position Rv3899c^136-157^ showed the motif to have a possible bifunctional NADP phosphatase/NAD kinase role (E-value: 0.37). Additionally, the domain of unknown function 5631 was also found to be conserved in motif 4 (Rv3899c^297-346^) while DUF5632 was conserved in motif 2 (Rv3899c^193-226^) and 5, respectively.

**Figure 2:**
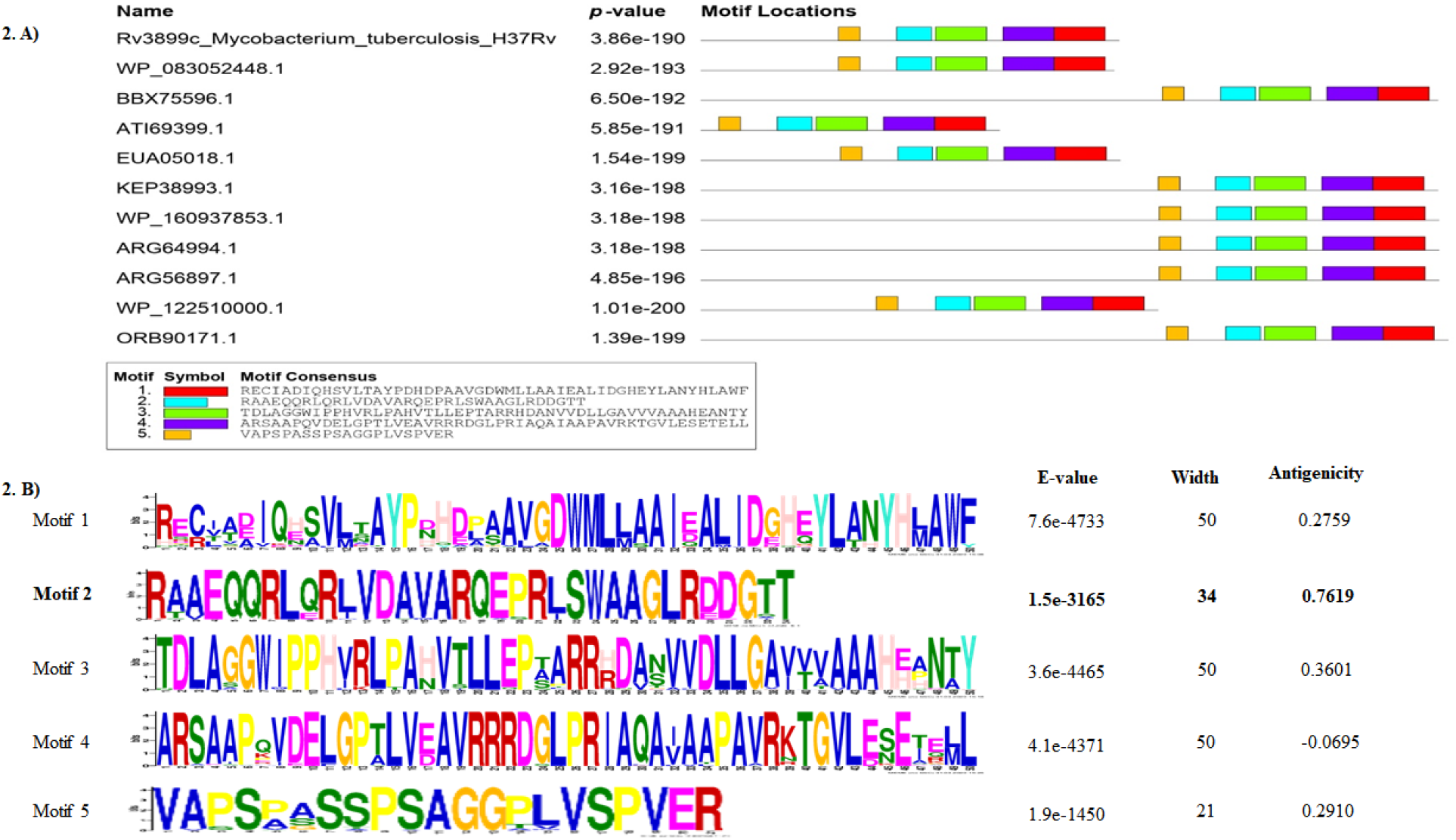
**A)** Occurrence of overlapping motifs in Rv3899c and related homologs. **B)** Motif sequence analysis shows ‘motif 2’ (RAAEQQRLQRIVDAVARQEPRISWAAGLRDDGTT) to have high antigenicity among the predicted motifs.

### 3.3. Physio-chemical characterization of Rv3899c protein of H37Rv

Rv3899c protein has 410 amino acids out of which 8.04 % are negatively charged (Asp+Glu) and 6.58 % are positively charged residues (Arg+Lys). The protein has a GRAVY score of 0.134 and instability index of 50.41 which classifies it to be a hydrophobic protein. A high aliphatic index of 88.49 predicts the protein to be possibly thermostable too. Overall, the protein shows weak polarity and high levels of hydrophobicity (Figure 3: A and B). Additionally, the protein has low flexibility and fairly accessible regions (Figure 3: C and D). The secondary structure of the protein as predicted by ‘PredictProtein Program’ indicates the protein to predominantly have 83.7% random coils (and 16.3% alpha-helix) and the extended strands were predicted to be altogether absent. The protein is also predicted to have Protein kinase C phosphorylation site at 191, 254, 296, 399 residues and Casein kinase II phosphorylation site at 13, 152, 249, 310 residues and N-myristoylation site at 4, 17, 50, 66, 164, 184, 267, 337, 348, 403 residues respectively.

**Figure 3:**
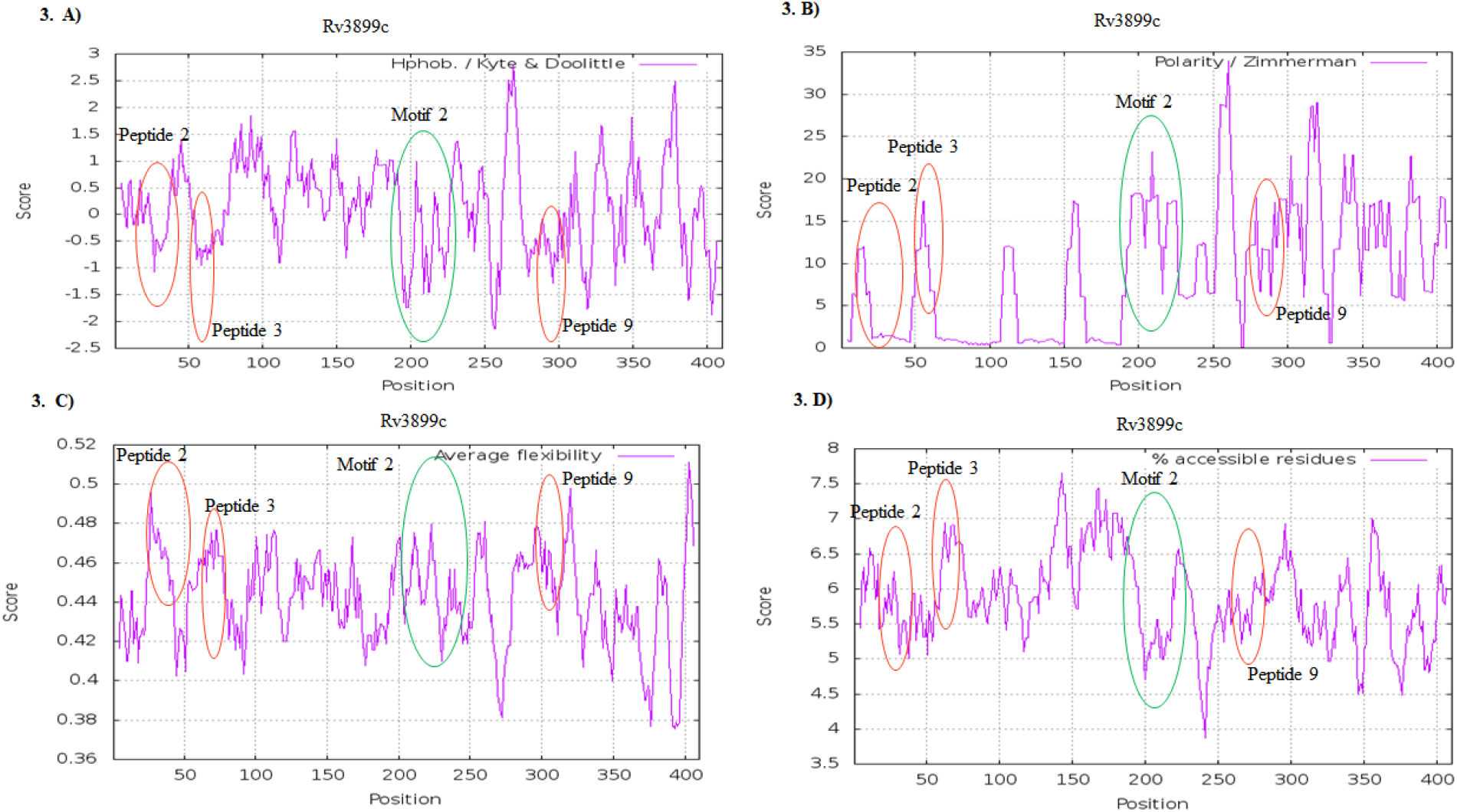
Profiles obtained in the analysis of the primary structure of Rv3899c. **A)** Hydrophobicity **B)** Polarity **C)** Average Flexibility and **D)** Flexibility of Rv3899c residues. Highlighted in a green empty circle is ‘motif 2’ and red empty circles are peptides 2, 3 and 9. These regions are the hydrophilic accessible residues in Rv3899c predicted to have high antigenicity.

### 3.4. Fold pattern and interaction network of Rv3899c

The fold pattern in Rv3899c is an ‘immunoglobulin-like’ fold in the sequence.Protein-protein interactions studies were done by STRING analysis, which predicted interaction of Rv3899c with ten other proteins (Rv2633c, Rv3896c, Rv0023, Rv3900c, eccB2, eccC2, eccD2, eccE2, PPE26 and PPE65) from H37Rv based on the fundamentals of gene fusion, co-occurrence, co-expression and gene neighborhood analysis. Rv3899c was predicted to interact with ESX-2 secretion system proteins eccB2, eccC2, eccD2 and eccE2 based on gene neighborhood and co-occurrence and co-expression in H37Rv. On the basis of gene fusion, gene neighborhood analysis and gene co-occurrence in H37Rv, Rv3899c appears to interact with Rv3990c and Rv3896c, which are poorly characterized alanine-rich proteins, and with Rv0023, a possible transcription regulator (Figure 4).

**Figure 4:**
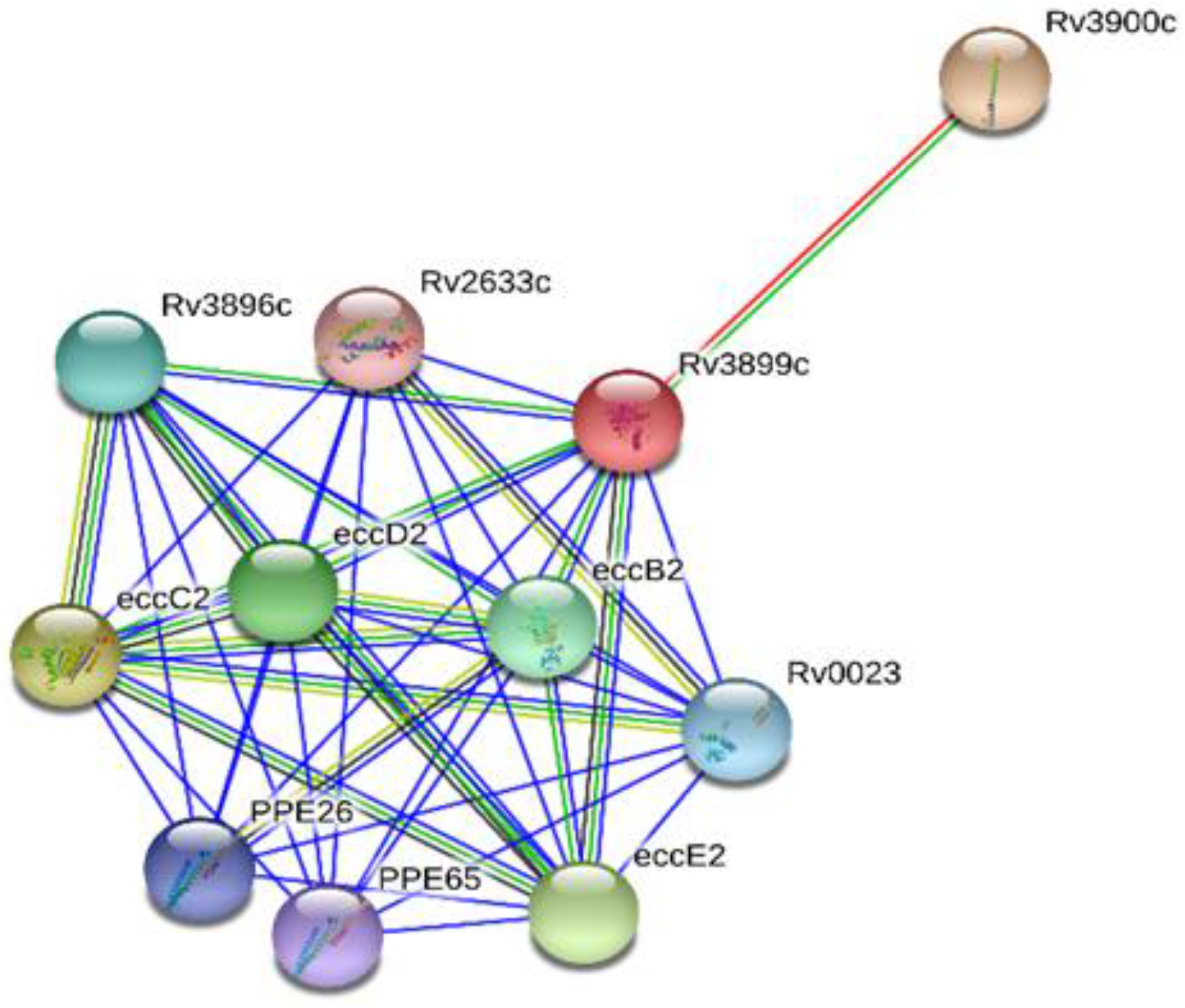
Protein-protein interaction network of Rv3899c showing PPE26 and PPE65 as the functional immunogenic partners and interaction with ESX-2 secretory system proteins.

### 3.5. Prediction of subcellular localization and toxicity of Rv3899c

Rv3899c protein was found to be secretory in nature based on various *ab initio* predictions on post-translational and localization aspects of the protein. Based on sequence property and peptide composition, PSLpred suggested the protein to be extracellular in nature, though a signal peptide wasn’t located in the protein and TMHMM and HMMTOP tool indicated absence of transmembrane helices as well.The non-toxic nature of Rv3899c was determined by BTXpred.

### 3.6. Characterization of antigenic determinants in Rv3899c

Rv3899c was found to have an antigenicity score of 0.5441. On sequence analysis, 10 antigenic peptides (Table 1) and several B-cell epitope rich segments were found with an AAR of 29.2 and 34.1 (Table 2). A high antigenicity score of > 1.0 was predicted for the peptides 5 and 7, which exceeds the VaxiJen tool standard threshold of 0.4. Peptide 10 also showed high antigenicity score (0.85) and was evaluated to have a protein-peptide binding site. Additionally, ‘motif 2’ (RAAEQQRLQRIVDAVARQEPRISWAAGLRDDGTT), a conserved region in Rv3899c and related homologs were predicted to be well accessible (Figure 3: D) with potentially high antigenicity score (0.76) and to contain a peptide-binding site. Apart from ‘motif 2’, three antigenic peptides (2, 3 and 9) were assessed to be predominantly polar using the GRAVY index. BLASTp analysis did not show the protein to have any similarity with human proteins. A comparison of Rv3899c with other experimentally validated antigens from H37Rv was done. On the basis of SecretomeP and antigenicity score, all the antigens were predicted to be secretory against a standard threshold of 0.5. Rv3899c and its predicted interacting partner Rv3621c (PPE65) had the highest SecP score of 0.950344 and 0.956296, respectively and antigen Rv3418c (groS) had the lowest (0.630609) (Table 4). VaxiJen v2.0 analysis predicted Rv3874 (cfp-10) to constitute the highest antigenicity score (0.7826) among the antigens studied followed by Ag85B with a score of 0.5727. Overall, antigenicity score of Rv3899c (0.5441) was found to be similar or somewhat higher than that of Rv1790 (0.5049), Rv1860 (0.5233), Ag85A (0.5259), Ag85C (0.4402), Rv3418c (0.5170) and Rv3621c (0.52) (Table 3).

**Table 1:**
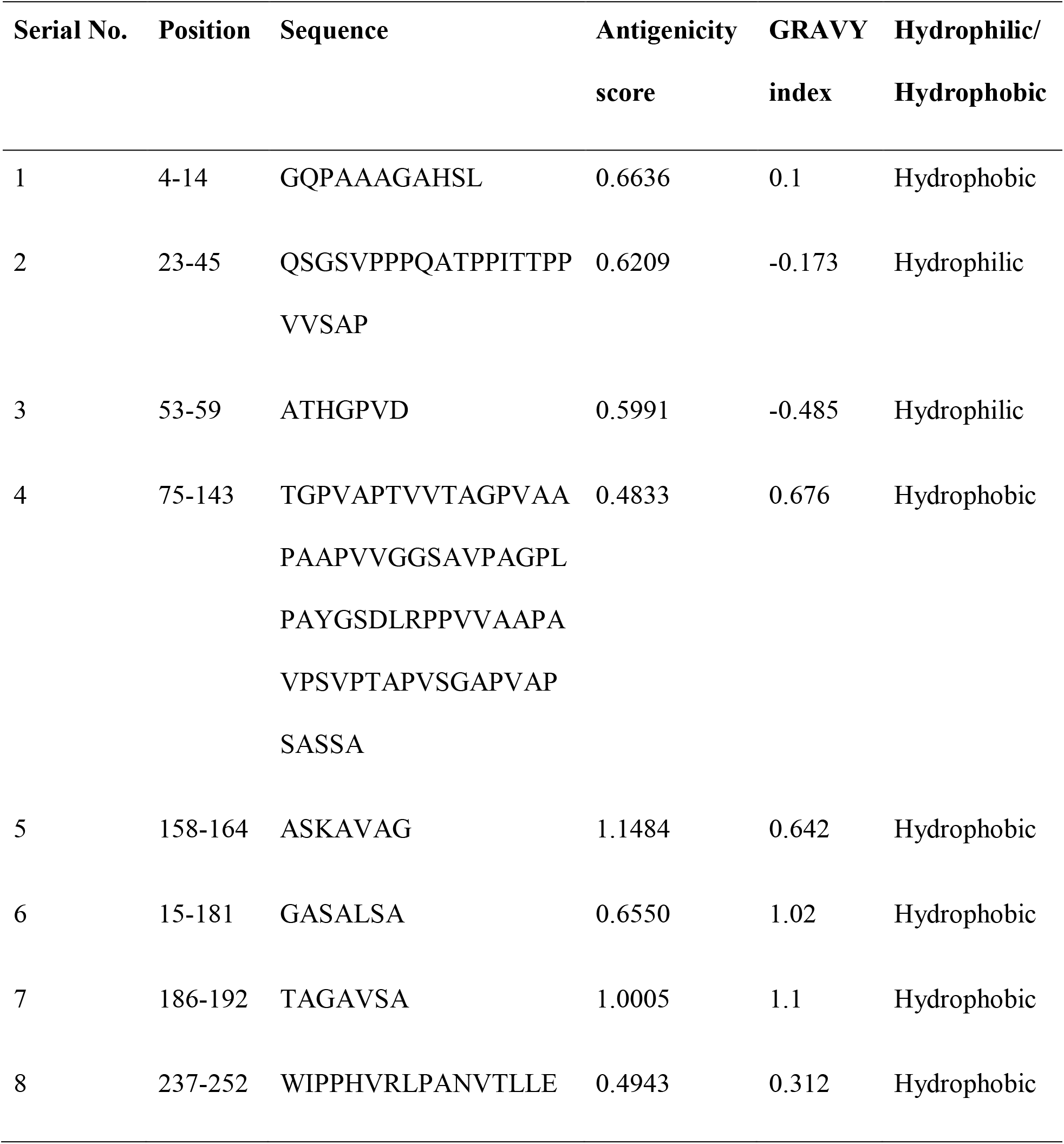
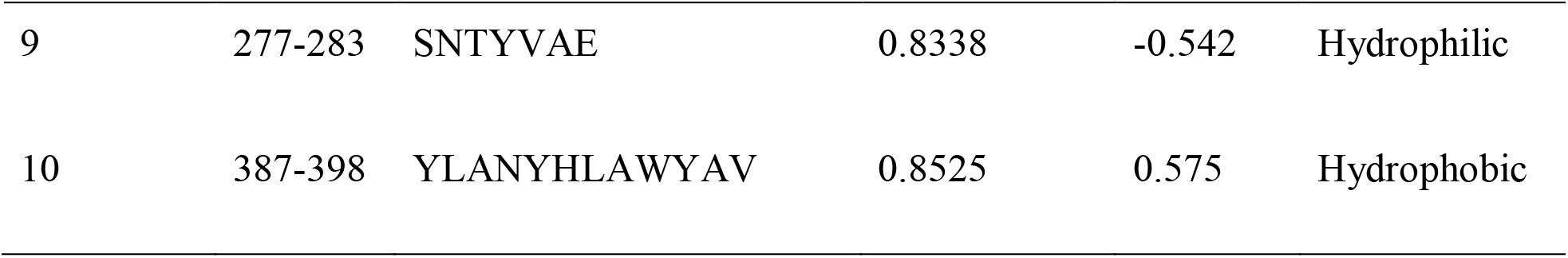
Prediction of antigenic segments in Rv3899c using Immunomedicine tool and analysis of their antigenicity using VaxiJen v2.0 and polarity profile using ProtScale of ExPASy.

**Table 2:**
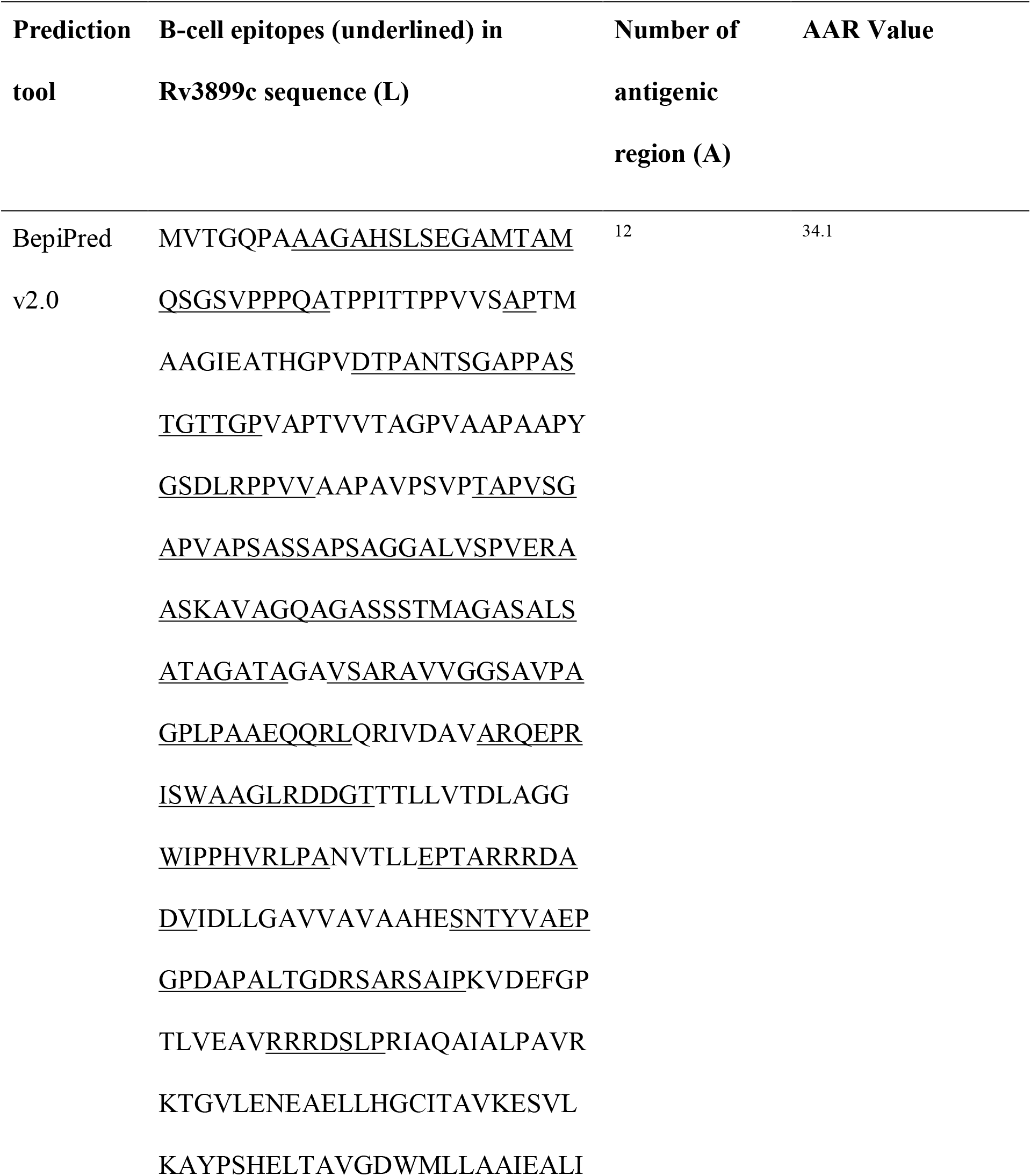
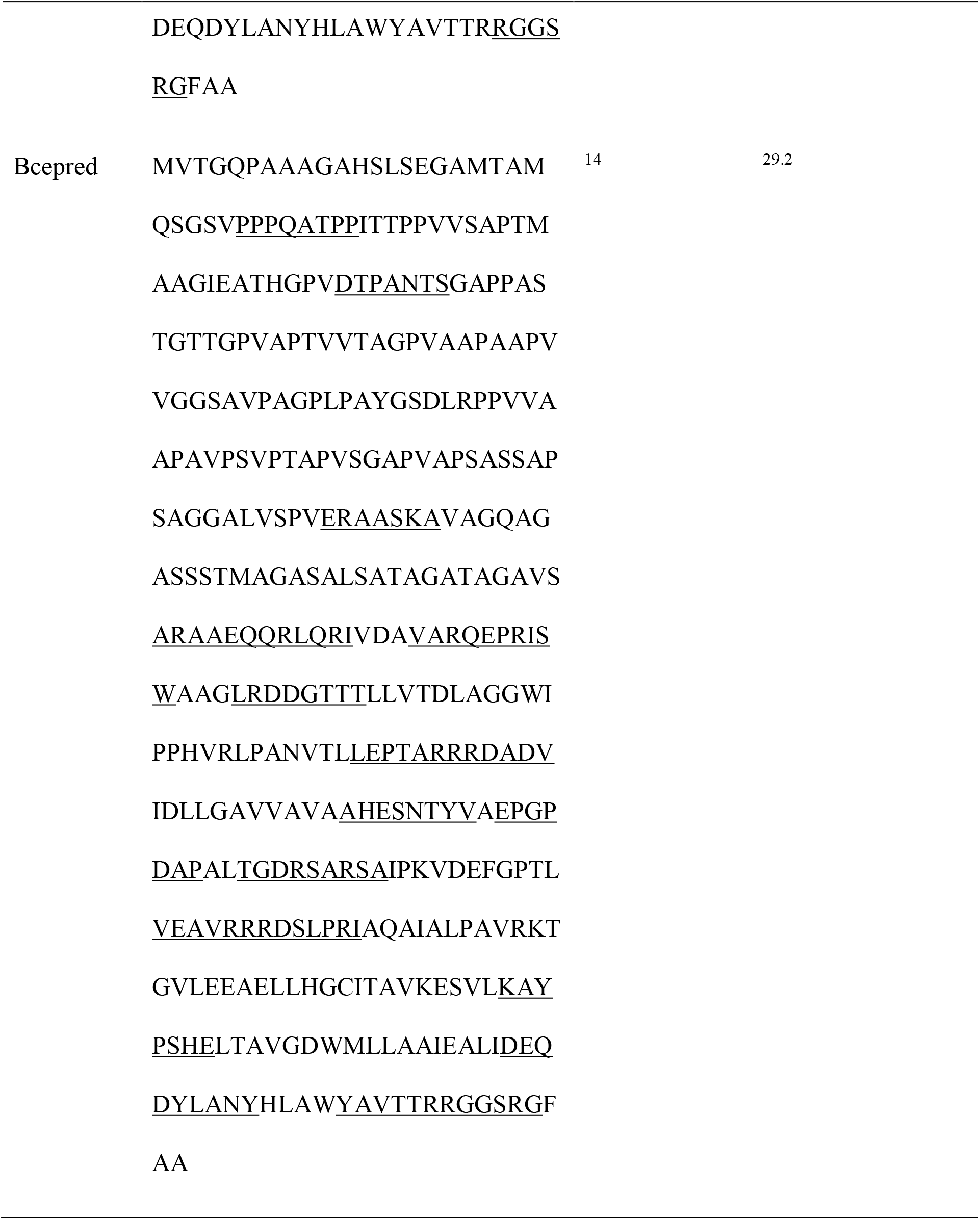
Analysis of B-cell epitope rich regions in Rv3899c and calculation of relative Abundance of Antigenic Regions (AAR).

**Table 3:**
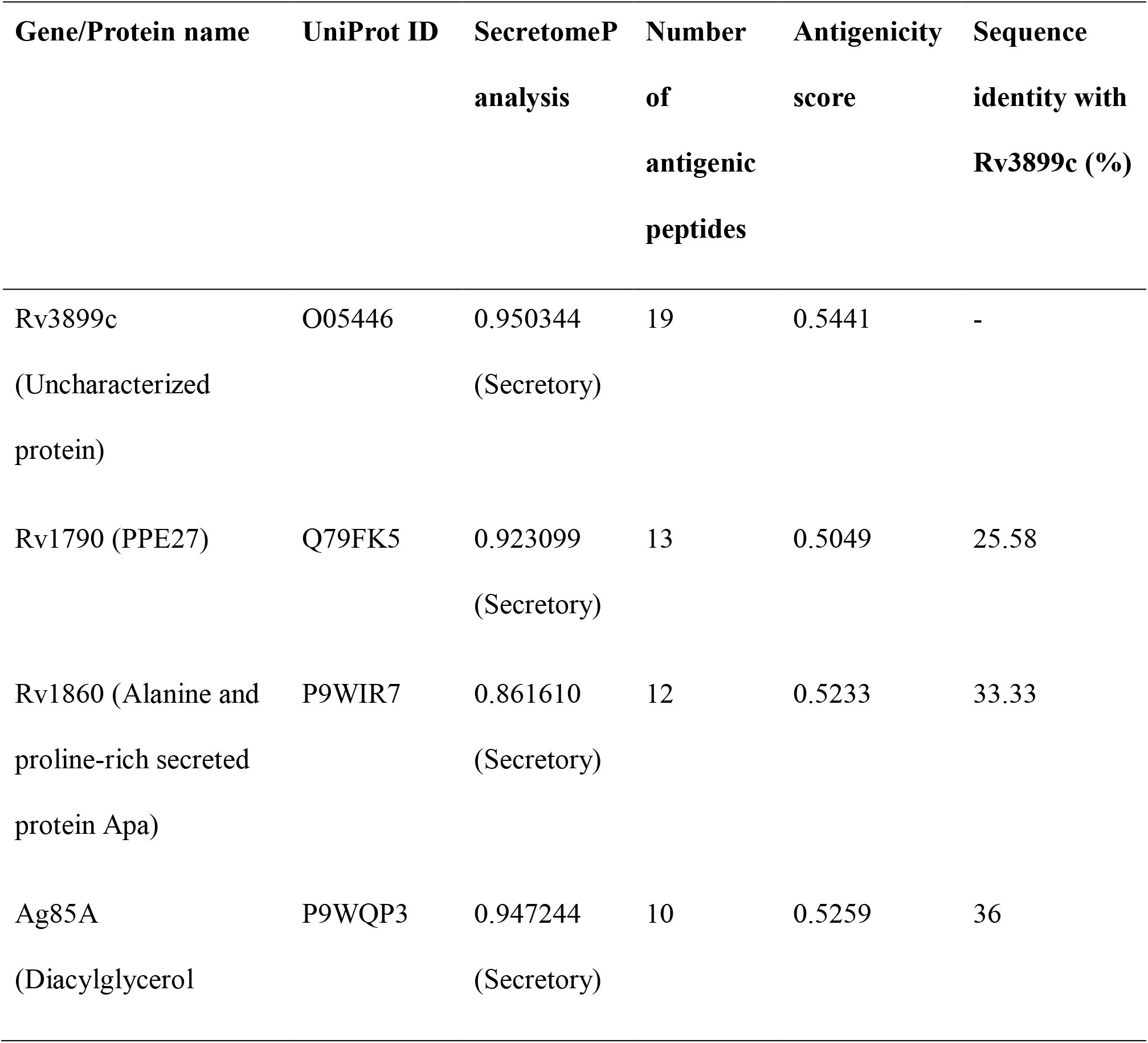
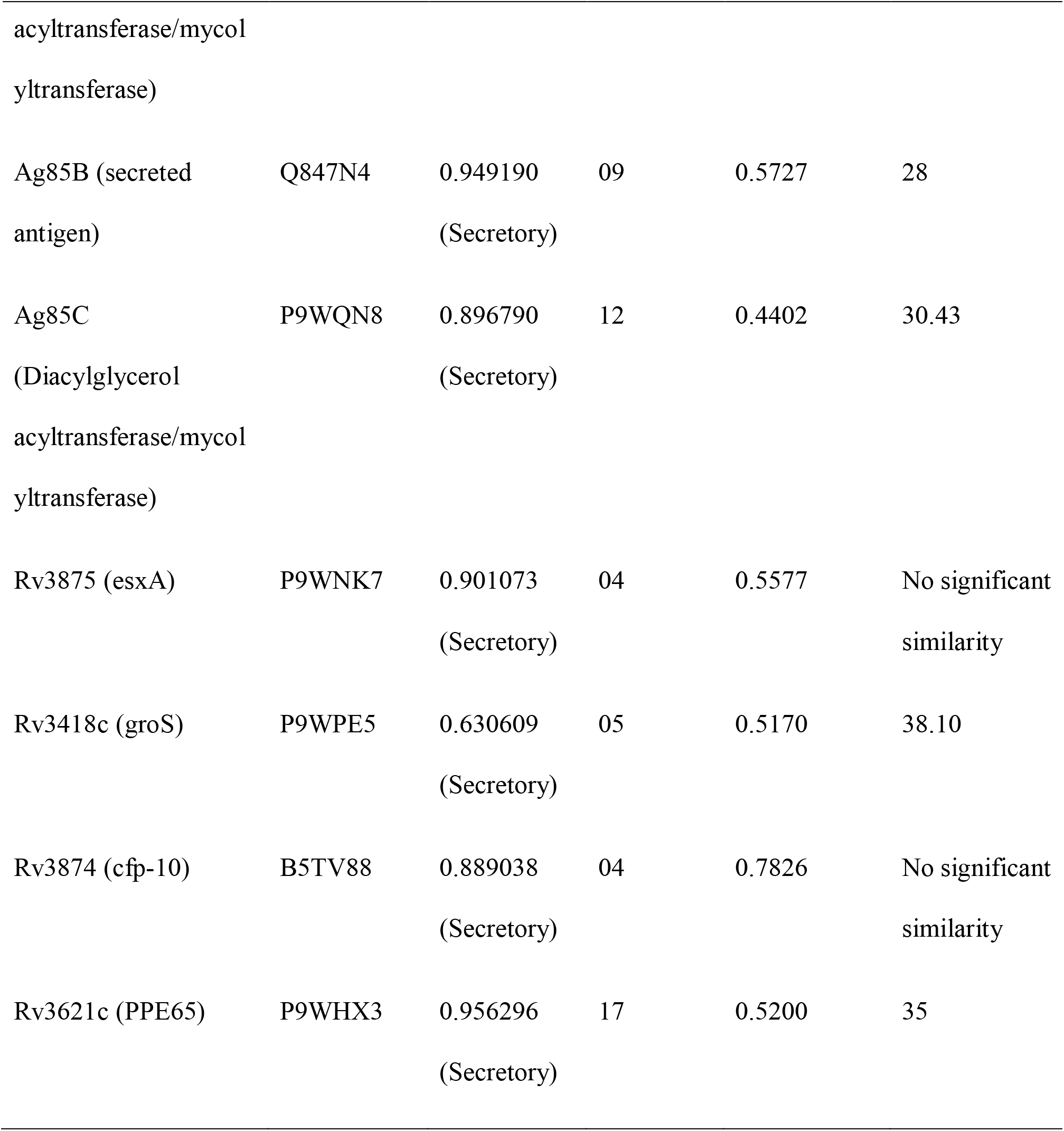
Comparative analysis of Rv3899c with other known mycobacterial antigens on the basis of antigenicity and SecP score.

PepBank alignment analysis predicted an antigenic peptide (ATHGPVDTPAN) in Rv3899c with a B-cell epitopeto show sequence similarity with a membrane protein peptide (GTMGPVWTPGN) from *Legionella pneumophila* that has been experimental proven to exhibit cross-reactivity with polyclonal anti-serum [53].

### 3.7. Mapping ‘motif 2’ in the secondary and tertiary structure of Rv3899c^184-410^

‘motif 2’ (RAAEQQRLQRIVDAVARQEPRISWAAGLRDDGTT), an antigenically conserved homolog rich in B-cell epitope, was mapped on the secondary (Figure S3) and tertiary structure (5IMU) (Figure 5. A) of Rv3899c^184-410^ procured from the PDB database.

**Figure 5:**
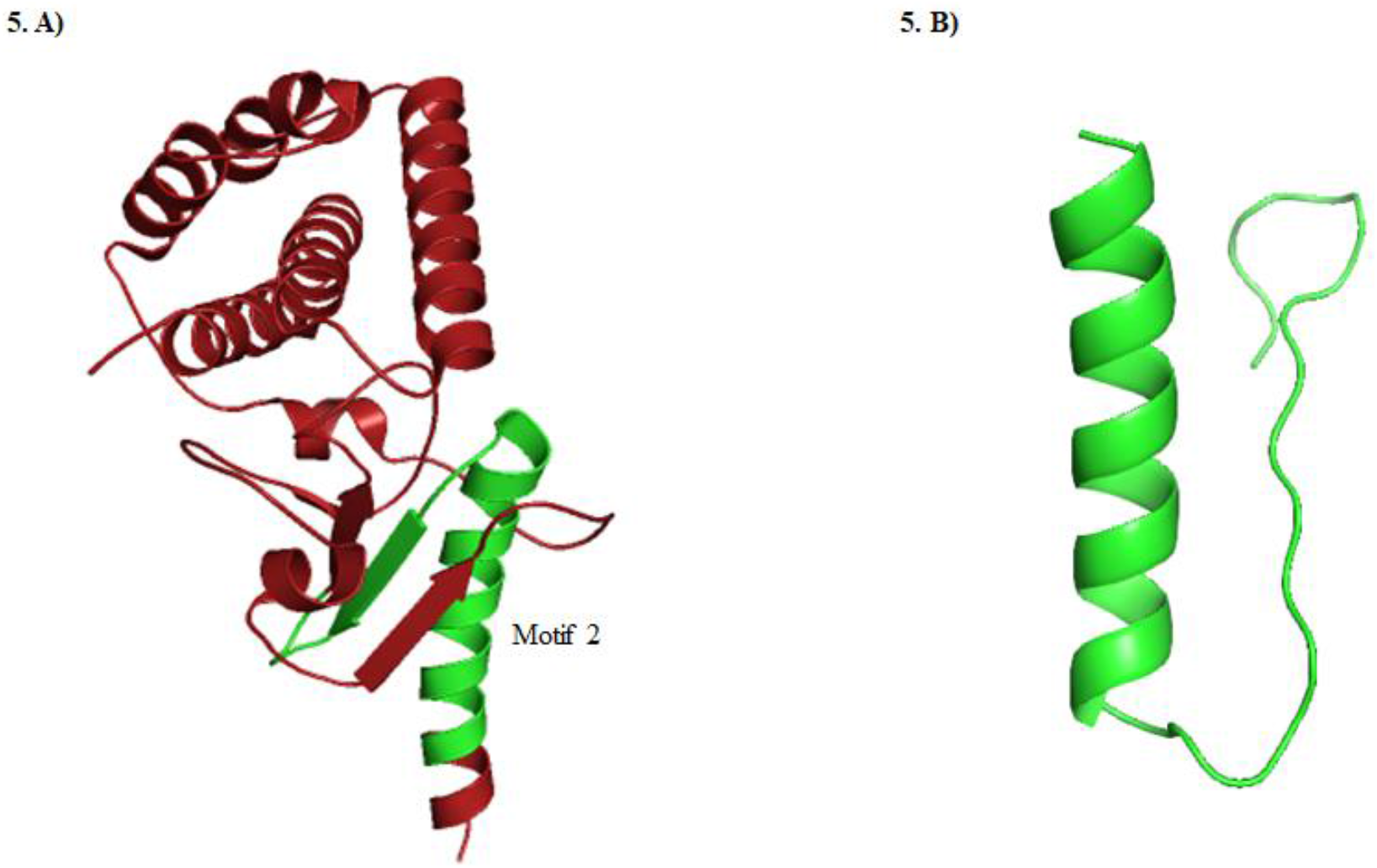
**A)** Rv3899c^184-410^ 3D structure ribbon model highlighted in red. Highlighted in green is the predicted antigenic conserved homolog ‘motif 2’ ‘193-226’ in the protein with B-cell epitope and protein-peptide binding site. **B)** 3D structure of conserved antigenic ‘motif 2’ (RAAEQQRLQRIVDAVARQEPRISWAAGLRDDGTT) built using SWISS-MODEL tool.

### 3.8. Citrullination and molecular docking of ‘motif 2’ with HLA-DRB1*04:01

T-cell prediction tools predicted the presence of T-cell epitopes for HLA-DRB1*04:01 in ‘motif 2’. Hence, ‘motif 2’ was separately modelled (Figure 5. B) from the crystal structure (5IMU) of Rv3899c^184-410^ and its binding affinity for HLA-DRB1*04:01 was evaluated using PatchDock server. The tool predicted binding energy of −88.68 kcal/mol and a docking score of 10570 (Figure 6) which indicates motif 2 as a good interacting partner of HLA-DRB1*04:01.

**Figure 6:**
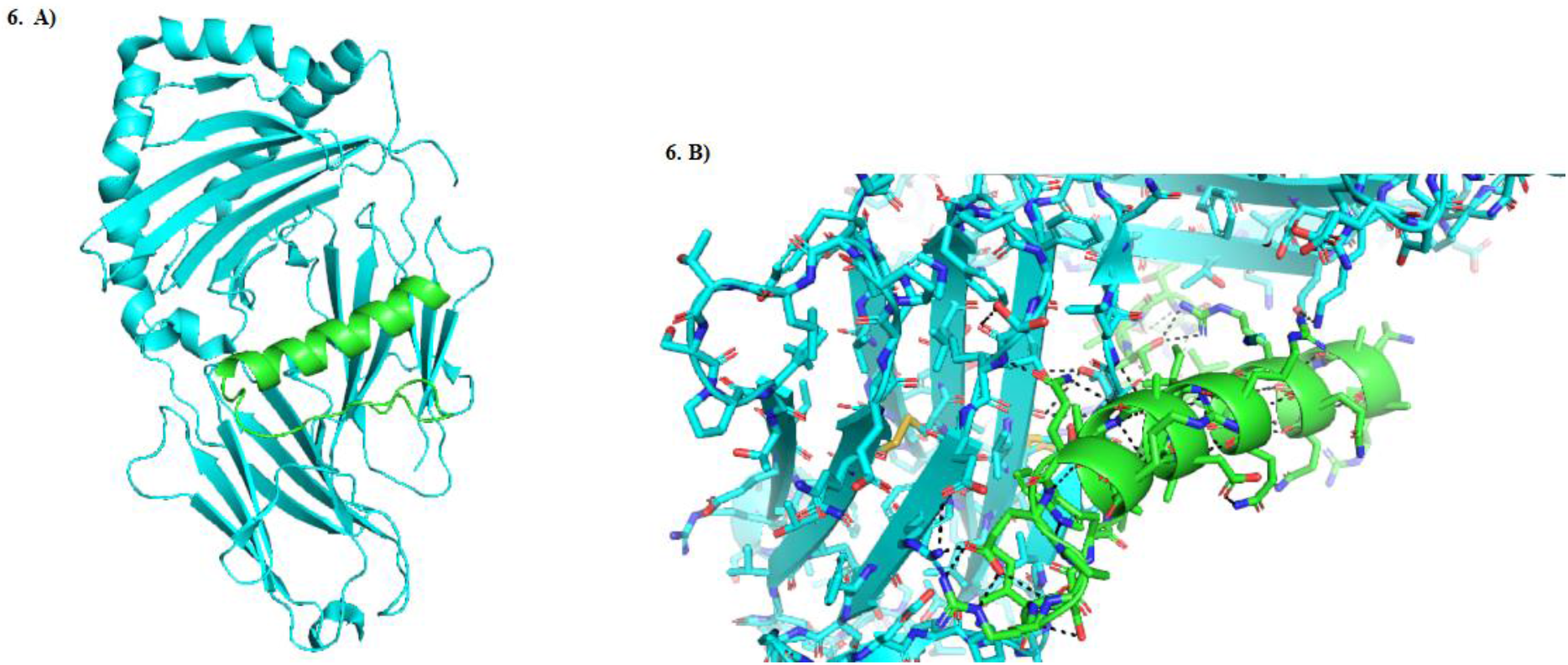
Molecular docking of HLA-DRB1*04:01 with ‘motif 2’. **A)** Helix-loop-helix structure of docked HLA-DRB1*04:01 and ‘motif 2’. **B)** Bird’s eye view of HLA-DRB1*04:01 (in blue) shows interaction (in black) with ‘motif 2’ (in green) with a binding affinity of −88.68 kcal/mol.

Citrullination of Arg1 of ‘motif 2’ was carried out using PyTMs plugin of PyMOL which can introduce common post-translational modifications in protein and peptide models. Docking analysis of the citrullinated peptide with HLA-DRB1*04:01 predicted enhanced binding energy of −193 kcal/mol and a docking score of 11830 as compared to the native peptide.

## 4. Discussion

The evolution of drug-resistant TB is a global threat that needs urgent intervention. With the emergence of clinical isolates of *Mycobacterium tuberculosis* with multiple or total resistance to the available TB drugs, our current methods of dealing with this infectious disease remain inadequate and the important requirement of an effective vaccine is strongly felt [54]. Development of immunotherapeutic vaccines generally includes mycobacterial antigenic proteins or peptides, live or attenuated *M. tuberculosis* or its whole-cell extracts [55]. Secretory antigenic proteins of H37Rv are frequently studied for their role as effective sub-unit vaccines and several of them such as ESAT-6, MTSA-10 and Rv1860 has been shown to elicit a protective immune response by activating mast cells, macrophages and dendritic cells against TB infections [56, 9]. Therefore, the screening and characterization of other mycobacterial secretory proteins with comparable or higher antigenicity can help us identify and investigate a new vaccine candidate. In this study, the hypothetical secretory protein Rv3899c of H37Rv has been extensively characterized and studied for its antigenic potential by using several computational tools. Interestingly, we found a Duffy-like binding domain of *Plasmodium* merozoites and bifunctional NADP phosphatase/NAD kinase role in the discovered motifs of Rv3899c. A Duffy-like binding domain is found in Duffy binding proteins in *Plasmodium vivax* and *Plasmodium knowlesi* merozoites that invade human erythrocytes. These proteins have been the key targets to develop *Plasmodium vivax* vaccines [57]. While the significance of the presence of a Duffy binding domain in Rv3899c is not understood, it indicates the possible immunogenic role of this protein, which needs to be ascertained through *in vitro* experiments. Also, Rv3899c was found to have an ‘immunoglobulin-like’ fold in the sequence. Proteins consisting of an Ig-like fold usually interact with other Ig-like domains and have a role in immune function [58].

STRING analysis was performed to study the protein-protein interaction of Rv3899c. The tool predicted the protein to interact with four ESX-2 secretion system proteins (eccB2, eccC2, eccD2 and eccE2) which are membrane proteins that help in the transportation of secretory antigens across the cytoplasm [59]. This interaction of Rv3899c with the ESX-2 secretion system proteins indicates its possible secretory pathway and antigenic role. The antigenic PE/PPE proteins of H37Rv (PPE26 and PPE65) were also predicted to be the interacting partners of Rv3899c. PPE65 has been experimentally shown to induce the stimulation of IFN-γ in bovine cell lines while the immunogen PPE26 has been shown to instigate both innate and adaptive immune response in humans by interacting with TLR2 which leads to the activation of mitogen-activated protein kinase (MAPK) and nuclear factor kappa B (NF-kappa-B) signaling pathways. It also stimulates the activation of macrophages through the increase of pro-inflammatory cytokine production by stimulating the expression of cell surface molecules such as CD80 and CD86 [45].

To estimate the antigenic potential of Rv3899c, a comparison of its antigenicity score was done with the score of the known immunogenic proteins from Mtb, using the VaxiJen tool that assigns an antigenicity score based on physicochemical characterization. We found Rv3899c’s antigenicity score (0.54) to be at par with ESAT-6 (0.56) which has been reported to induce a protective immune response in mice [8] and with Rv1860 (0.52), which enhances the proliferation of IFN-γ from healthy latently infected human CD4+ and CD8+ T-cells [9] Significantly, antigenic peptides 5, 7 in Rv3899c were predicted to have an antigenicity score >1, higher than 0.7 of ‘motif 2’, which is a conserved domain across several mycobacteria and its antigenicity score is better than most of the known immunogens. On further analysis, it was also observed that these antigenic determinants are well accessible in the protein and do not show similarity with human proteome.

To characterize the antigenicity of Rv3899c, tools for the prediction of B-cell epitopes were used. For predicting B-cell epitopes, two different algorithms are applied: Bcepred, which predicts linear continuous epitopes in antigenic sequences using physicochemical properties [34] and BepiPred-2.0, which is based on a random forest algorithm [35]. The Abundance of Antigenic Regions (AAR) was then determined to normalize the results predicted from these two tools. Rv3899c was found to have an AAR of 29.2 and 34.1, respectively which is comparable with the average AAR of Excretory/Secretory (ES) proteins from *M. tuberculosis* H37Rv (40.63) and *M. tuberculosis* Beijing isolate 48 (37.55) [60], hence indicating it to have strong antigenic potential.

The human leukocyte antigen complex (HLA) is a dense and polymorphic region present on the chromosome 6p21.3. The class II of the HLA genes are responsible for expressing cell-surface glycoproteins on dendritic cells, macrophages and antigen-presenting cells. An important role of these glycoproteins is to introduce peptides to the T-cell receptors before the activation of T-cells [49]. Several reports have also demonstrated the association of the alleles of HLA class II with pulmonary TB [61]. In this study, we wanted to understand whether ‘motif 2’ from Rv3899c interacted with HLA-DRB1*04:01, the HLA class II allele against which the T-cell epitope was predicted. Docking studies predicted a binding affinity of −88.68 kcal/mol. Interestingly, previous studies on peptide interaction with HLA-DRB1*04:01 have demonstrated that substitution of arginine to citrulline at specific surface positions allows for high-affinity binding. This happened due to the transformation of the imino group to an uncharged carbonyl group, which showed enhanced peptide-MHC affinity and elicited CD4+ T cell response in mice [51]. To find out whether a similar effect can be observed in the ‘motif 2’ of Rv3899c, we performed *in silico* citrullination of Arg1 followed by docking, which predicted binding energy of −193 kcal/mol. Although there are six Arginine residues in ‘motif 2’, we carried out citrullination of Arg1 because it was found to be in the accessible region of the protein (Arg193). A decrease in the binding score indicates the important role citrullination of Arg1 could play in the binding of ‘motif 2’ to HLA and further substantiates the probable use of this peptide inRv3899c in developing targeted vaccines for TB infections.

With the increasing number of drug-resistant TB infections, it is important for us to unravel the function of various encoded proteins of *M. tuberculosis.* However, 27% of its proteome is still uncharacterized [62]. An attempt has been made in this paper to predict the function and antigenicity of the hypothetical protein Rv3899c and also to identify its interacting partners. On the basis of our analysis and characterization, Rv3899c makes for a strong candidate immunogen and we believe the computationally predicted immunogenicity with a potential to confer defence against TB infections needs to be validated experimentally.

## 5. Conclusion

To summarize, Rv3899c of theoretical molecular mass 40.7 kDa is a secretory hypothetical protein with 10 highly antigenic determinants and a potential vaccine candidate for TB. The non-toxic protein without a homolog in humans consists of ten interacting partners which include ESX-2 system proteins (eccB2, eccC2, eccD2 and eccE2) and mycobacterial known antigens (PPE26 and PPE65). Additionally, ‘motif 2’, a conserved hydrophilic region in Rv3899c and related homologs was predicted to be a B-cell and T-cell epitope rich antigenic region and to have an affinity for HLA-DRB1*04:01. Collectively, results from this study provide insights into the secretory Rv3899c protein of *Mycobacterium tuberculosis* (H37Rv) that could be evaluated further for next-generation TB control approaches.

## Acknowledgements

We thank the Principal, Acharya Narendra Dev College, University of Delhi for the infrastructural support and research facilities.

## Availability of data and material

Data available within the article or its supplementary materials.

## Code availability

Not applicable

