## Supplemental File for "Computational identification and characterization of antigenic properties of Rv3899c of *Mycobacterium tuberculosis* and its interaction with Human leukocyte antigen (HLA)"

**Address for communication:**

Dr. Urmi Bajpai (Associate Professor)
Department of Biomedical Science,
Acharya Narendra Dev College (University of Delhi),
Govindpuri, Kalkaji, New-Delhi-110019,
India.
Contact Number: +91-9811299719
Fax Number: +91-11-26412547

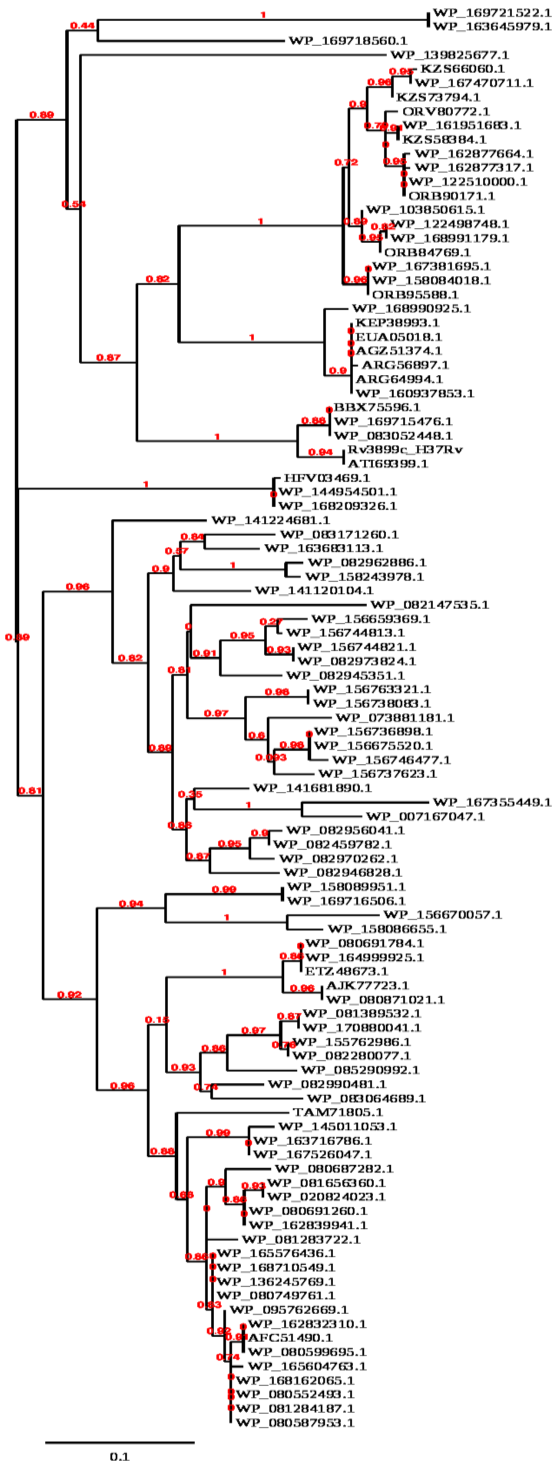


**Figure S1:** Analysis of homology of Rv3899c of H37Rv with other microorganisms through Phylogenetic tree analysis.

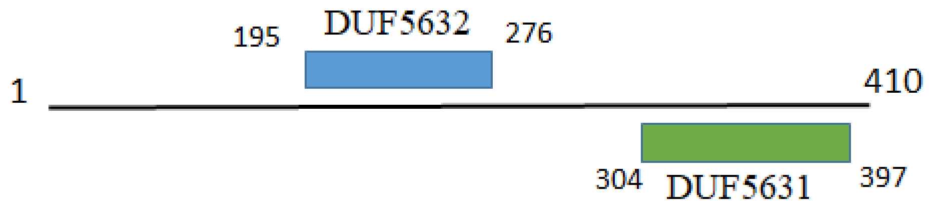


**Figure S2:** Schematic representation of the domain architecture of Rv3899c. Two domain of unknown function DUF5631 (195-276 aa) and DUF5632 (304-397 aa) are present.

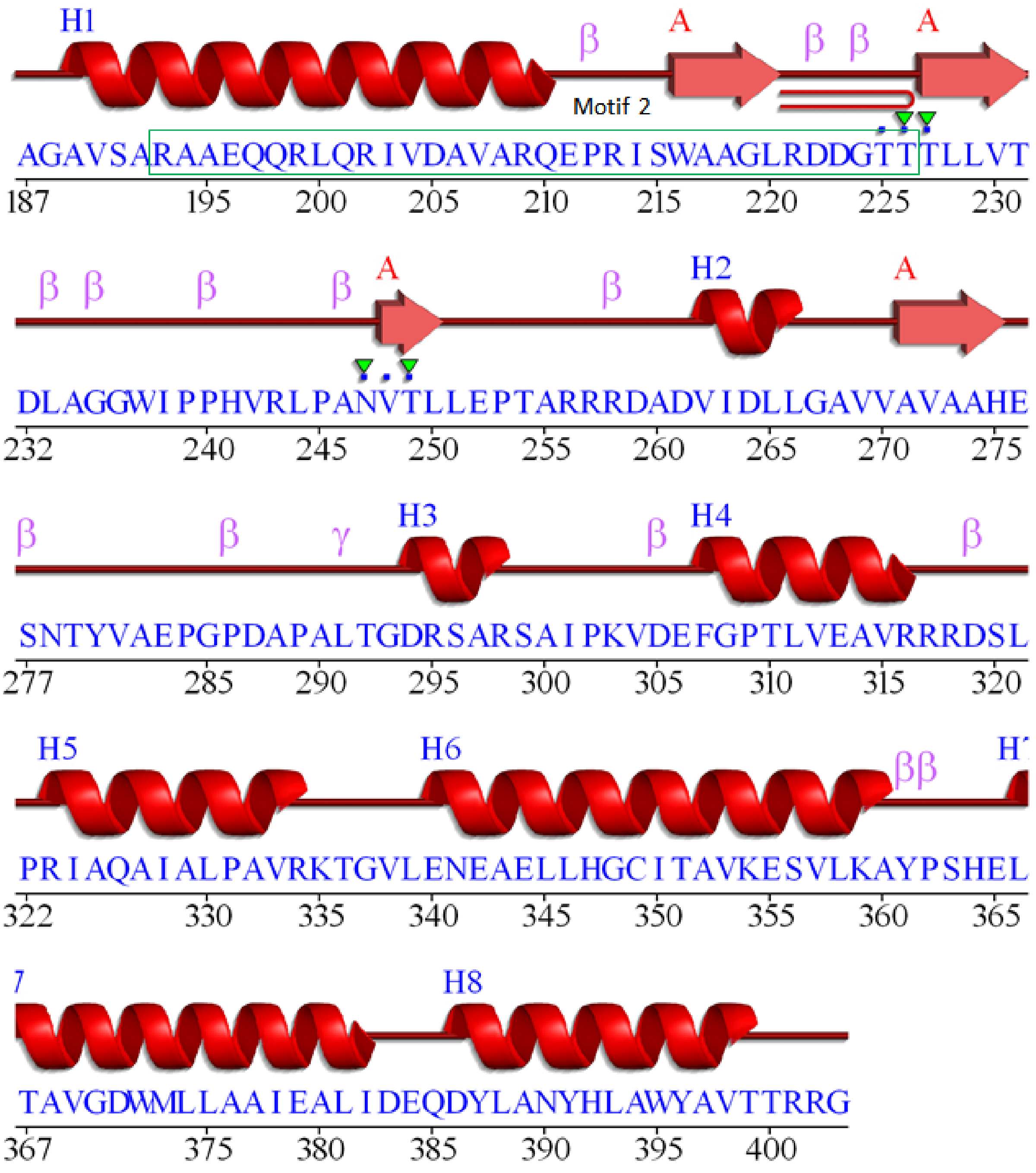


**Figure S3**: Secondary structure of Rv3899c$^{187\text{-}410}$. The region marked by a green box is the predicted antigenic conserved homolog 'motif 2'. H: Helix, A: Chain name, β: beta-turn and γ: gamma turn. A hairpin loop was predicted at site ‘221-226’ and residue contact site to metal was discovered at sites ‘225-227’ and ‘247-249’, respectively.
